# Spontaneous variability predicts adaptive motor response in vocal pitch control

**DOI:** 10.1101/2020.06.06.138263

**Authors:** Ryosuke O. Tachibana, Mingdi Xu, Ryu-ichiro Hashimoto, Fumitaka Homae, Kazuo Okanoya

## Abstract

Our motor system uses sensory feedback to keep behavioral performance in desired status. From this view, motor fluctuation is not simply ‘noise’ inevitably caused in the nervous system, but should provide a role in generating variations to explore better outcomes via their sensory feedback. Vocal control system offers a good model to investigate such adaptive sensory-motor interactions. The pitch, or fundamental frequency (FF), of voice is adaptively regulated by hearing its auditory feedback to compensate FF deviations. Animal studies, particularly for songbirds, have demonstrated that the variability in vocal features contributes to the adaptive control, although the same issue in human vocalizations has remained unclear. Here, we tested whether and how the motor variability contributes to adaptive control of vocal FF in humans. We measured the amount of compensatory vocal responses against FF shifts in the auditory feedback, and quantified the motor variability as amplitudes of spontaneous FF fluctuations during no shift vocalizations. The result showed a positive correlation between the ratio of compensation and the spontaneous vocal variability. Further analysis indicated that this correlation was due to slowly fluctuating components (<5 Hz) of the variability, but not fast fluctuations (6-30 Hz), which is likely to reflect controllability from the central nervous system. Moreover, the compensatory responses consisted of the same frequency range with the slow component in the spontaneous variability. These findings consistently demonstrated that the spontaneous motor variability predicts the adaptive control in vocal FF, supporting the motor exploration hypothesis.

**Significance statement:** We regulate our own vocalization by hearing own voice. This fact is typically observed as canceling-out (compensatory) responses in vocalized pitch when artificial pitch shifts were induced in the auditory feedback of own voice. Interestingly, the amount of such compensation widely ranges among talkers from perfect cancellation to almost nothing. Here we demonstrated that participants who spontaneously exhibited larger fluctuations showed greater amounts of the compensation against feedbacked pitch shifts. Our in-depth analyses showed that slowly fluctuating components in spontaneous pitch variability are specifically correlated with the compensation ratios, and was shared in the compensatory response as a dominant component. These findings support the idea that such variability contributes to generating motor explorations to find better outcomes in motor controls.

## Introduction

Precise control of vocal pitch, or fundamental frequency (FF), is essential for human communication since the vocal FF is a dominant cue for prosodies in speaking, or melodies in singing. A key aspect of the vocal control is hearing own voice, or the auditory feedback. Speakers regulate their own vocal FF by canceling out subtle FF deviations induced in the auditory feedback (Elman, 1981; Kawahara, 1994; Burnett et al., 1998; Larson et al., 2000). For example, shifting up vocal FF in the auditory feedback elicits a response shifting down FF in the vocalization. Such compensatory vocal response does not always cancel out the shift completely, but rather remains around half or less of the induced shift with large individual differences (Hain fet al., 2000; Liu and Larson, 2007; Liu et al., 2010; Scheerer and Jones, 2012). Investigating mechanisms underlying the compensatory responses for vocal FF regulation provides opportunities to understand the adaptive audio-vocal system, which plays a critical role in our vocal control.

Recent studies in animal vocalizations, particularly in birdsongs, have suggested that variability in vocal features contributes to vocal adaptation against errors induced in the auditory feedback (Tumer and Brainard, 2007; Sober and Brainard, 2012; Kuebrich and Sober, 2015; Woolley and Kao, 2015; Tachibana et al., 2017). Songbirds typically vocalize stereotypic songs in adulthood that have almost identical acoustical patterns across renditions, while exhibiting slight but unignorable variations in their acoustical features such as FF. These variations have been reported to contribute to maintaining the song quality (Kao et al., 2005; Tumer and Brainard, 2007; Charlesworth et al., 2011). In particular, the FF shifts in the auditory feedback elicit compensative responses of vocal FFs in birds’ song syllables (Sober and Brainard, 2009). The amount of this compensation became larger when distributions of original and shifted FF variations are more overlapped (Sober and Brainard, 2012; Kuebrich and Sober, 2015), linking the wider variability with the greater vocal adaptations. It has also been shown that temporal patterns of FF fluctuation within a brief sound element guide to keep and improve the song quality (Charlesworth et al., 2011; Kojima et al., 2018). Intriguingly, the vocal variability in birdsongs is not simply due to the intrinsic noise in the peripheral motor system, but a certain amount of them is ‘actively’ generated by a dedicated circuit that is required for song learning (Kao and Brainard, 2006; Hampton et al., 2009; Olveczky and Gardner, 2011; Kojima et al., 2018). These findings in songbirds’ vocalization have supported the idea that motor variations contribute to adaptive controls by generating the motor exploration (Wu et al., 2014; Woolley and Kao, 2015; Dhawale et al., 2017). Moreover, the active generation of variability in the motor processes is likely to suit to the adaptation-related motor exploration (Dhawale et al., 2017). Such mechanism for songbirds’ vocal control could be shared with humans (Hahnloser and Narula, 2017), especially when taking into account behavioral and neural parallels between these two species for vocalization development (Doupe and Kuhl, 1999; Kuhl, 2004; Lipkind et al., 2013; Tchernichovski and Marcus, 2014; Prather et al., 2017).

In contrast, relationships between variability and adaptability in human vocal control have not been well documented. Variability in the human vocal FF appears to consist of several components reflecting different sources or mechanisms. These components have been classified according to their dominant frequencies in the modulation spectrum, which is an amplitude spectrum of FF changing frequency (modulation frequency). For example, a quasi-periodic FF fluctuation during singing (or *vibrato*) has been reported to show a peak around 4–7 Hz on the modulation spectrum, with greater stability in trained singers (Sundberg, 1987; Shipp et al., 1988; Howes et al., 2004). In contrast, non-periodic components at relatively higher modulation frequencies at 10–20 Hz, or fine fluctuation (Akagi et al., 1998; Akagi and Kitakaze, 2000; Saitou et al., 2005), have been reported to be involved in the perception of voice quality both in speaking (Akagi et al., 1998) and singing (Akagi and Kitakaze, 2000). Such aperiodic fast fluctuation is likely due to the physiological instability of peripheral vocal organs (Schoentgen, 2002), and hence, is less or not controllable for the central nervous system. These reports lead to a question of whether and to what extent these different types of variability could contribute to the vocal regulation.

Here, we assessed associations between vocal compensatory responses against auditory feedback modifications and variabilities of different components in vocal FF trajectories to obtain a better understanding of how we accomplish adaptive vocal regulations based on the auditory feedback. In the experiment, the vocal FF in the auditory feedback was modified while participants were vocalizing, and the rate of compensation in their vocalized FF was measured. We quantified the vocal variability that was spontaneously generated in vocalizations for unmodified feedback after separating the variability components into different modulation frequency bands. By correlation analyses between the variability and the compensation ratio, we found a greater correlation in slowly fluctuating components than fast fluctuations that are likely to be less controllable in the central nervous system. Further analysis showed that the compensatory response consists of the frequency range of the slow component in the spontaneous fluctuation. These results consistently support the hypothesis that the spontaneous variability subserves motor explorations to enhance the compensatory response against perturbations in the auditory feedback.

## Results

### Variety of the compensation ratio across participants

In the experiment, participants were asked to continuously produce isolated vowels for two seconds twice while listening to auditory feedback via headphones, and only the second voice was modified in its feedback (**Fig. 1A**; see Methods for detail). We found a clear tendency of compensation (canceling out) in vocalized FF against the artificially induced FF shifts in auditory feedback (**Fig. 1B**). The amount of compensation was almost proportional to the amount of seven FF shift conditions (0, ±25, ±50, or ±100 cents), as already shown in the previous study (Xu et al., 2020). Thus, we defined the compensation ratio for an individual participant as a sign-inverted slope of a fitted line to compensation amounts as a function of introduced FF shifts (**Fig. 1C**). The obtained compensation ratio was variable across participants with ranging from −0.13 to 0.82 (0.39 ± 0.21 [mean ± SD]; **Fig. 1D**).

**Figure 1.**
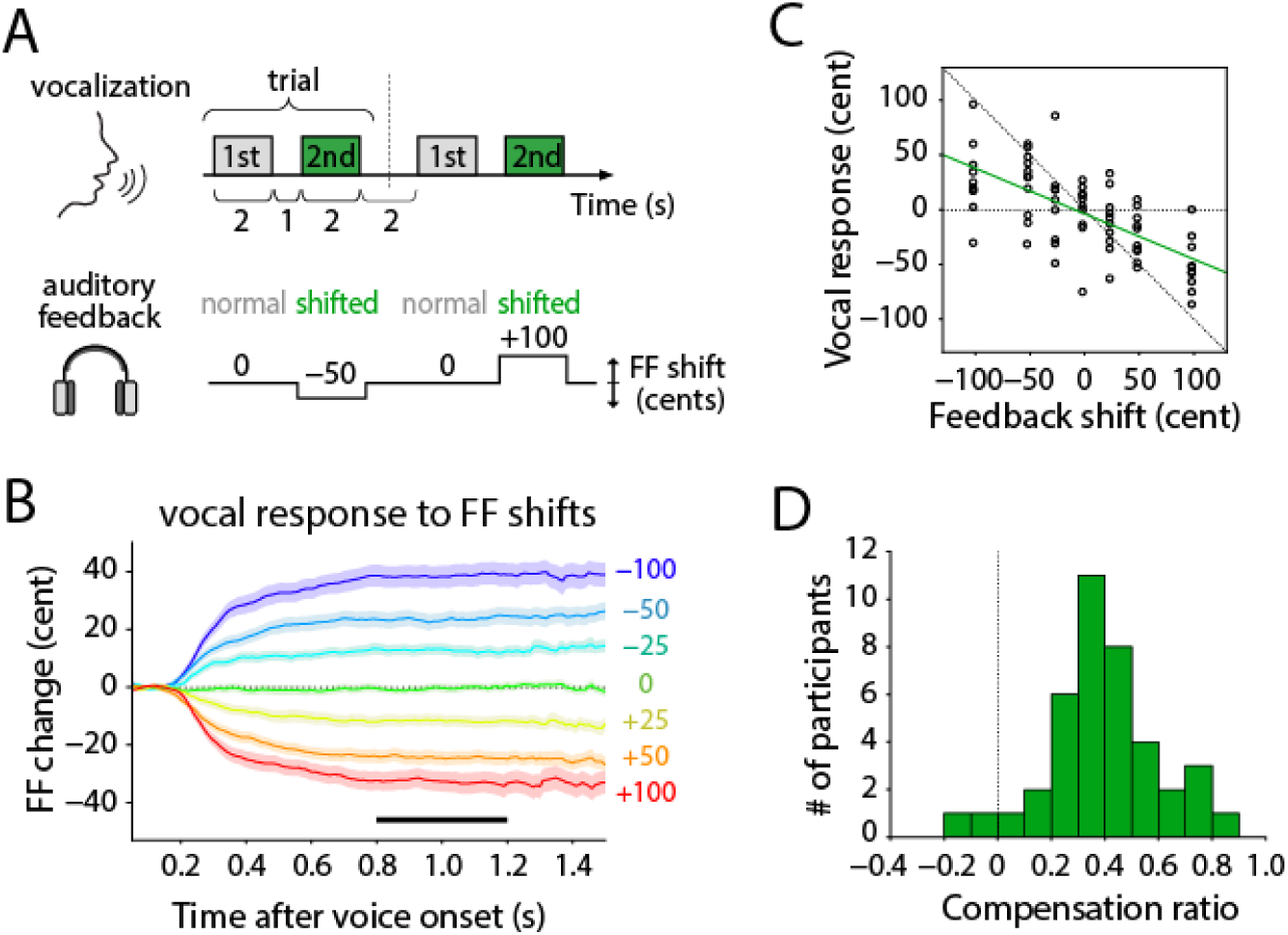
Measuring compensation responses in vocal fundamental frequency (FF) against artificially induced FF shifts in auditory feedback. **A**. Schematic drawing of the experimental design. Participants vocalized twice in one trial with normal auditory feedback for the first time, and with modified auditory feedback in the second time. **B**. Average of vocal FF change across all participants in response to seven conditions of the FF shift in auditory feedback (0, ±25, ±50, or ±100 cents). All trajectories were aligned at vocal onsets, and detrended before averaging (see Methods for detail). Pale-colored area indicates the standard error (*n* = 40). **C**. Example of compensation amounts as a function of FF shifts obtained from one participant (M04). Each dot indicates the compensation amount of each trial, which was calculated as an average of the plateau period (0.8–1.2 s after voice onset) indicated as a black bar in panel B. The compensation ratio was estimated as a sign-inverted value of the slope of fitted line, shown as a green line. Diagonal dotted line indicates sign-inverted unity slope **D**. Histogram of compensation ratios obtained from all participants.

### Variability in slow component of spontaneous fluctuations correlated with the compensation ratio

To assess what extent the motor variability related to the adaptation, we performed correlation analyses between the compensation ratio and several types of FF variability. Note that we only included participants who showed compensatory responses (i.e., positive value in the compensation ratio), resulted in excluding two out of forty participants from further analysis. To quantify vocal variability that was spontaneously generated without external perturbations, we calculated the standard deviation (SD) of an original FF trajectory of the first vocalization (no FF shift presented) in each trial. The mean of all SDs was defined as the variability of whole frequency components (“whole”). This variability ranged from 8.55 to 23.87 (14.19 ± 3.72) cents. We found the whole variability was significantly correlated with the compensation ratio (**Fig. 2A**; Spearman’s correlation coefficient *r*_s_ = 0.40, sample size *n* = 38, *p* = 0.014). Then, we aimed to divide the whole variability into slow or fast fluctuating components according to the modulation spectrum of the spontaneous FF fluctuation that was calculated by the 1/2-octave-band filter-bank method. The obtained modulation spectrum (**Fig. 2B**) showed apparent two peaks at modulation frequencies of 2–3 Hz and 6–10 Hz, suggesting two different variability components. None of the participants exhibited a sharp peak around 4–7 Hz corresponding to the presence of the vibrato component (Sundberg, 1987; Shipp et al., 1988; Howes et al., 2004). Thus, we defined slowly and rapidly changing components, termed as “slow” and “fast” fluctuations with having modulation frequency ranges of less than 5 Hz and 6–30 Hz, respectively (**Fig. 2C**). Obtained variabilities of slow and fluctuation components were ranged 7.99–22.52 (13.07 ± 3.72) and −2.04–6.93 (3.50 ± 3.72) cents, respectively.

**Figure 2.**
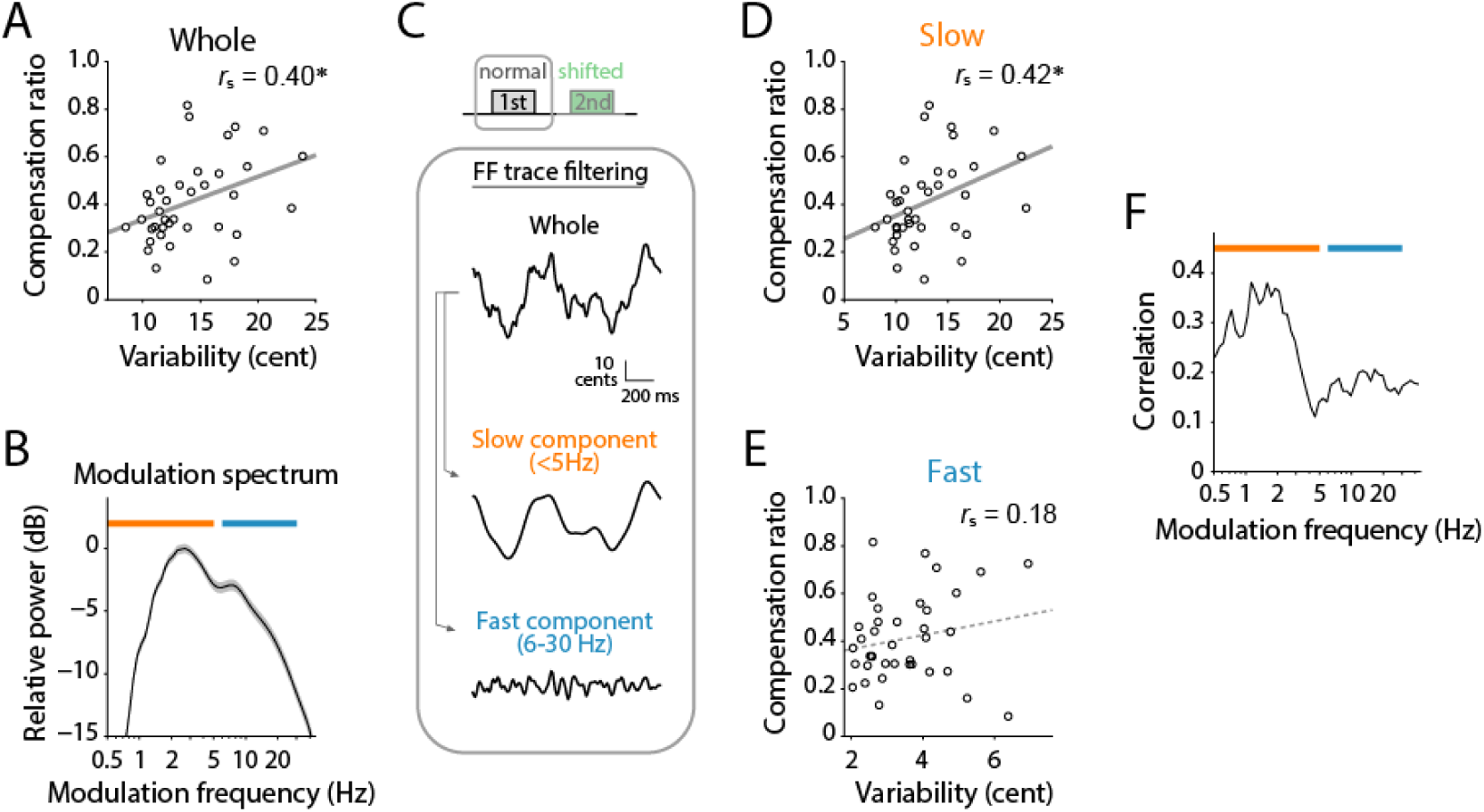
Spontaneous FF variability during vocalizations without modification in auditory feedback, and its relationship with the compensation ratio. **A**. The relationship between the compensation ratio and variability calculated from original (whole) FF trajectories during no FF shifts. Each circle indicates data from one participant. *r*_s_ shows Spearman’s signed-rank correlation coefficient. Two participants who showed negative values in the compensation ratio were excluded as outliers. Asterisk (*) indicates statistically significant correlation (*p* < 0.05). **B**. Modulation spectrum of spontaneous variation in vocal FF trajectories computed by a 1/2-octave filter bank. Gray area indicates the standard error among 40 participants. Orange and blue lines indicated frequency ranges of slow and fast fluctuation components. **C**. Examples of filtering on the original FF trajectory (whole) to obtain the slow and fast fluctuation components (slow: <5 Hz, fast: 6–30 Hz). **D,E**. Correlation between the compensation ratio and variability of slow (D) or fast (E) fluctuation components, respectively. **F**. Correlation coefficient (Spearman’s) between the compensation ratio and the variability of each modulation band as a function of center frequency of the half-octave filter bank.

The correlation analysis between these variabilities and the compensation ratio resulted in that the slow component showed a significant correlation (**Fig. 2D**; *r*_s_ = 0.42, *n* = 38, *p* = 0.009), whereas the fast component did not (**Fig. 2E**; *r*_s_ = 0.18, *n* = 38, *p* = 0.282). In addition to this result, the same tendency was observed in different vowels (see **Supporting Information**), providing further support for the finding that the larger slow component predicts the greater compensation. Moreover, to confirm the relative impact of each modulation frequency band on the compensation, we calculated the correlation coefficients between compensation ratios and variability values in each of the subbands which were derived from the modulation spectrum analysis. This analysis showed the consistent result (**Fig. 2F**) that the slow component (less than 4 Hz in modulation frequency) exhibited a greater correlation with the compensation ratio, but the rapid one (higher than 5 Hz) did not.

### Increase of slow component in compensatory response

To assess which frequency component in the FF trajectory the participants used to compensate for the FF shifts in auditory feedback, we compared variabilities in the second vocalizations (with FF shifts) with the first one (no shifts). We found significantly larger variability in ±100-cent shift conditions for the slow component (**Fig. 3A**; paired-t test, *t*(39) = −8.73, *p* < 0.001;) but not for the fast component (**Fig. 3C**; paired-t test, *t*(39) = −0.24, *p* = 0.814). The variability difference of the second from the first vocalization increased according to the amount of FF shift for the slow component (**Fig. 3B**), but remained constant around zero for the fast one (**Fig. 3D**). These results showed that the compensatory FF changes contain the same ranges in modulation frequencies with the slow component of spontaneously generated vocal variability (i.e., without FF shifts in auditory feedback). Further, we calculated the 2nd-1st variability difference in each of the subbands derived by the modulation filter bank to confirm the modulation frequency of the compensatory FF movement. The result (**Fig. 3E**) clearly depicted that the slow modulation component, which is associated with the compensation ratio in the spontaneous fluctuation (**Fig. 2F**), exhibited an extra variability for the compensatory vocal responses. This coincident finding strongly supported the idea that spontaneous variability in the slow components plays a critical role in the compensation.

**Figure 3.**
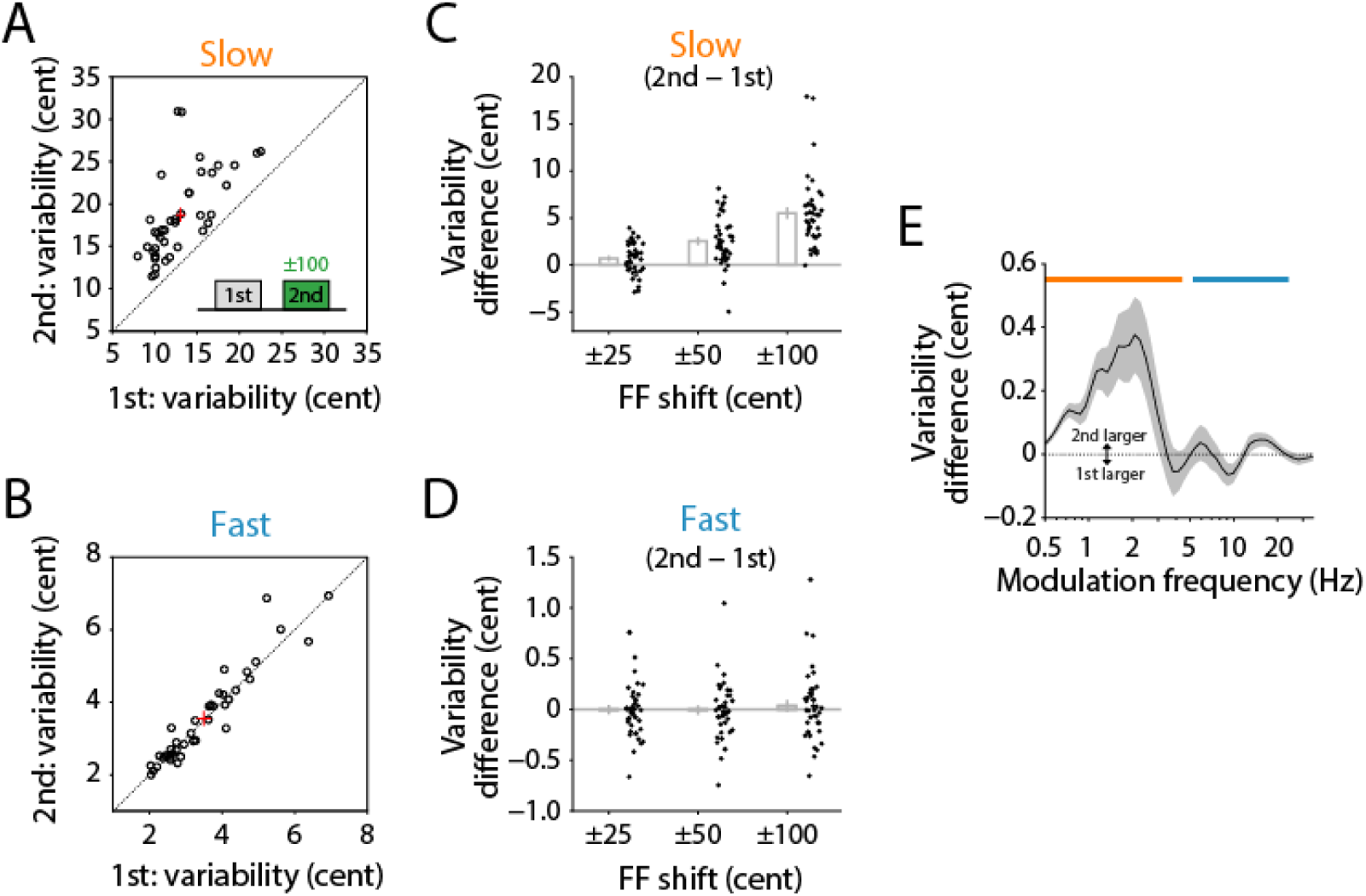
Variability comparison between the first (no FF shifts) and second (FF shifted) vocalization. **A,B**. Mean variability of the slow (A), and fast fluctuation (B) components in ±100-cent shift conditions of the second vocalization comparing to the first vocalization. Red crosshair indicates the mean and standard error. **C,D**. Variability difference of the slow (C) and fast (D) components in ±25-, ±50-, and ±100-cent shift conditions between the second and first vocalizations. Errorbar indicates the standard error among 40 participants. **E**. Variability difference of each subband component obtained by the modulation filter bank in ±100-cent shift conditions between the second and first vocalizations. Gray area indicates the standard error.

### Compensation ratio decreased with large FF shift

The motor exploration hypothesis predicts that the amount of compensation becomes small when the induced shift is large. For example, with a certain amount of variability, the originally intended FF will not be overlapped well with largely shifted versions of the FF distribution that reflects the motor exploration range (**Fig. 4A**). This can reduce opportunities to find correct (intended) FF during vocalization, and hence, decrease the compensation ratio for such large shifts. We tested this possibility by calculating the compensation ratio for each of the three shift amounts (**Fig. 4BC**). We pooled positive and negative shifts with inverting its sign. We statistically compared the compensation ratios among three conditions, and found significant difference between 50- and 100-cent shifts (**Fig. 4C**; Wilcoxon’s singed-rank test with Bonferroni correction; *z* = 3.48, *p* = 0.002), but not between 25- and 50-cent (*z* = −0.15, *p* = 1.000) or between 25- and 100-cent (*z* = 2.26, *p* = 0.072). While the compensation ratio in 100-cent shifts was significantly lower than others, its correlation with the variability of the slow component was still significant (**Fig. 4D**; *r*_s_ = 0.40, *n* = 38, *p* = 0.013). These results consistently supported the motor exploration hypothesis in vocal control.

**Figure 4.**
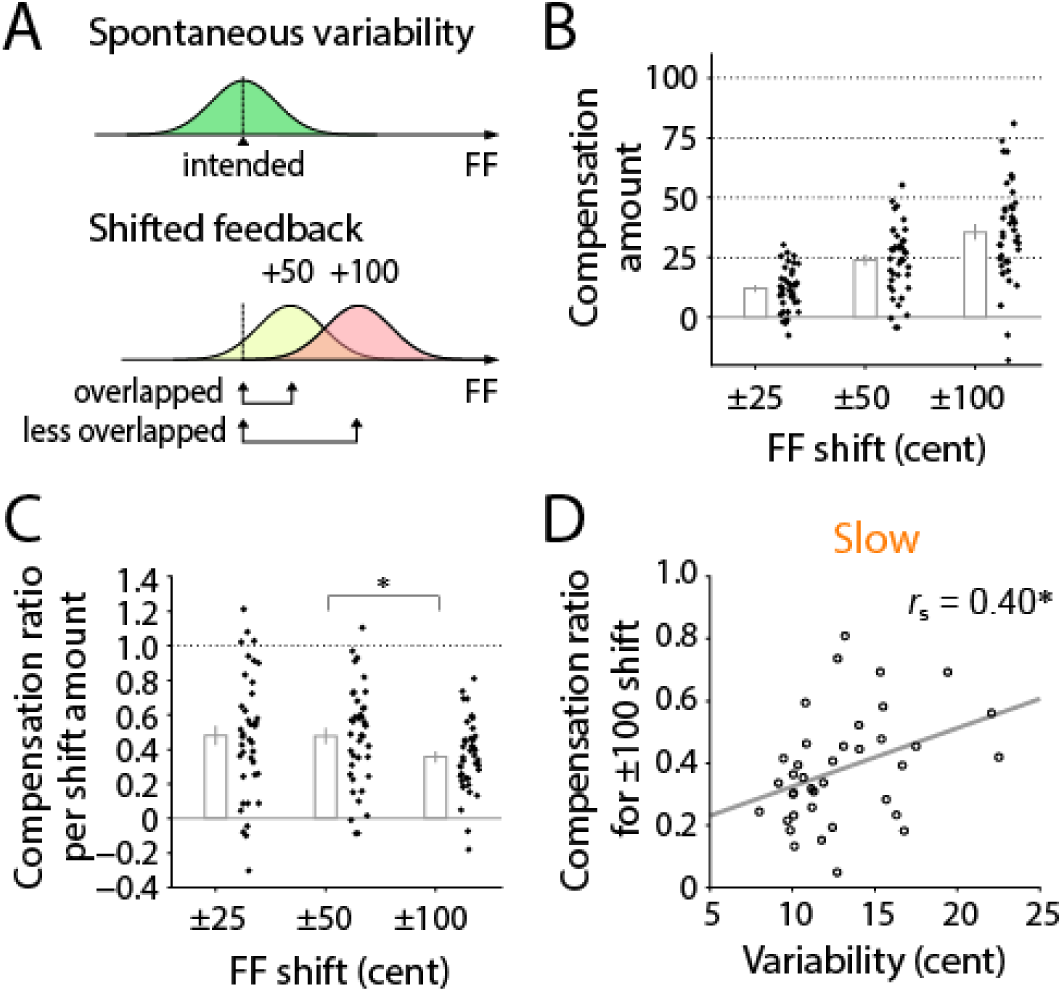
Decrease of compensation ratio for larger FF shifts. **A**. Schematic drawing of normalized distributions for the spontaneous FF variability (upper) and shifted versions of its feedback after the introduction of +50 and +100 cent shifts (lower). Given a certain amount of variability, originally intended FF will not be overlapped well with the distribution for large FF shifts, or will be outside of the motor exploration range. This can be expected to reduce the compensation ratio for that condition. **B**. Amount of compensatory responses against different amounts of FF shift (25, 50, and 100 cents). The vocal responses to positive FF shifts were sign-inverted and averaged with that to negative shift conditions. Each dot indicates individual participants. Error bar shows the standard error (*n* = 40). **C**. Compensation ratio obtained for each shift amount. The value was calculated as dividing the compensation amount (shown in A) by the shift amount (25, 50, or 100 cents). Asterisk (*) indicates statistically significant difference (*p* < 0.05; Bonferroni-corrected Wilcoxon’s signed-rank test). **D**. Correlation between the compensation ratio for the 100-cent shift amount and the variability of the slow component. *r*_s_ shows Spearman’s signed-rank correlation coefficient. Two participants who showed negative values in the compensation ratio were excluded as outliers. Asterisk (*) indicates statistically significant correlation (*p* < 0.05).

### Influence of perception and other factors

We additionally assessed other factors that potentially affect the compensation process, such as perceptual ability to discriminate vocal pitch. For this aim, we estimated participant’s ability to detect the FF shifts induced in recorded own voices using a dataset from the listening tests performed in our previous study (Xu et al., 2020). In this test, participants were asked to answer whether any pitch modification occurred in the second vocalization comparing with the first one in each trial (**Fig. 5A**). We estimated the discrimination threshold and accuracy for detecting the presence of pitch modification by fitting a sigmoid curve (**Fig. 5B**) on the detection rate dataset (see Method for details). Obtained discrimination thresholds and accuracies ranged 26.91–108.25 (54.71 ± 16.69) cents and 0.87–38.30 (14.13 ± 11.48) cents, respectively. We then tested correlations between these perceptual properties and the compensation ratio. The result showed that the compensation ratio did not significantly correlate with both the discrimination threshold (*r*_s_ = −0.17, *n* = 38, *p* = 0.298; **Fig. 5C**) or accuracy (*r*_s_ = 0.18, *n* = 38, *p* = 0.287; **Fig. 5D**), suggesting that the perceptual ability did not contribute the compensation in this case. Moreover, we tested if the amplitude of vocalization (or loudness level of auditory feedback) affected the compensation ratio. However, the relative amplitude level was not significantly correlated with the compensation ratio (*r*_s_ = 0.07, *n* = 38, *p* = 0.685; **Fig. S1D**). Lastly, we performed a stepwise multiple regression analysis to find the most effective model to explain the variation of the compensation ratio, amongst five explanatory variables: variability in slow and fast components, discrimination threshold, accuracy, and voice amplitude. The analysis best chose a statistical model that contained only the variability in slow component as an explanatory variable (adjusted *R*^2^ = 0.12, *df* = 36, *SSE* = 0.168, *p* = 0.019), indicating that the slow component is the main contributor for predicting the compensation ratio.

**Figure 5.**
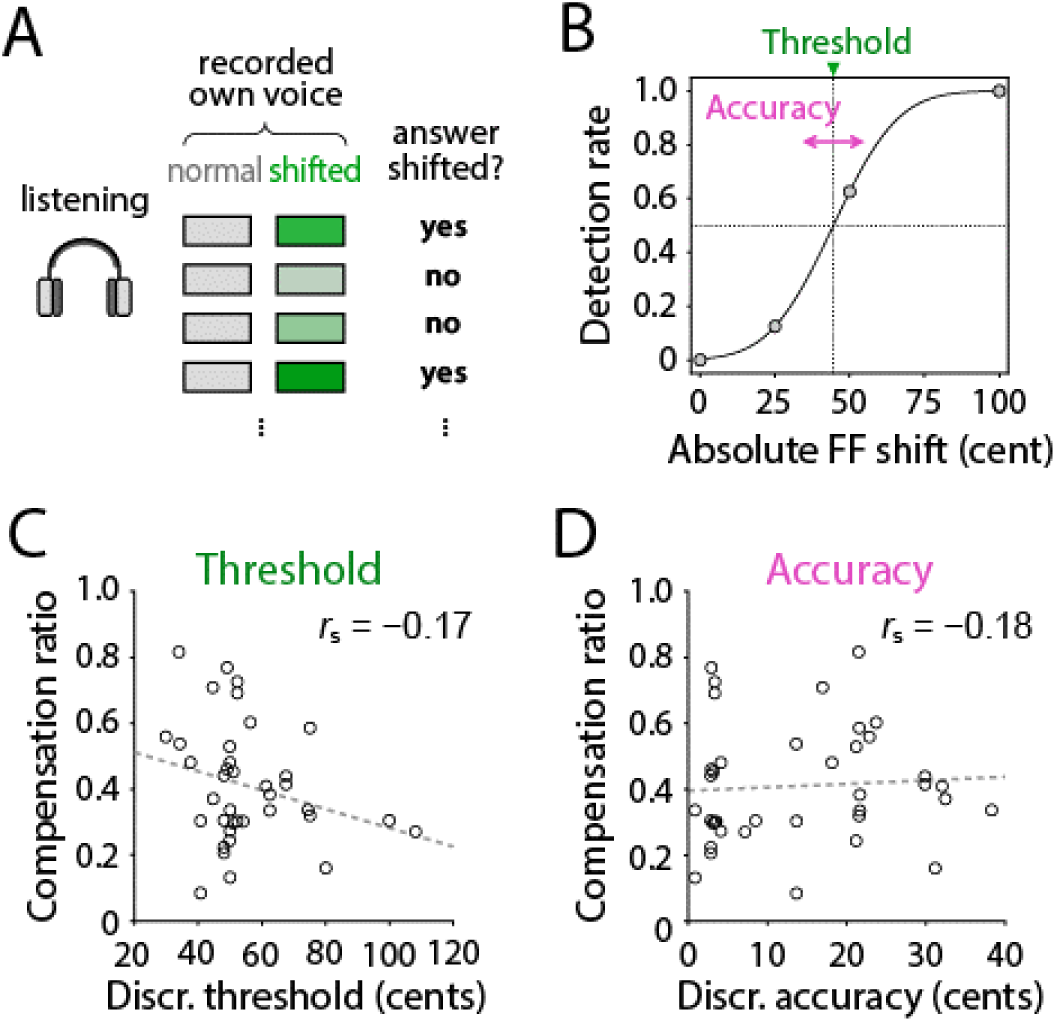
Participant’s ability to detect the FF shifts in recorded own voices, and its correlation with the compensation ratio. **A**. Test procedure. Participants listened to a pair of recorded voices corresponding the first and second vocalization in each of vocalization trials, and judged whether the second one had the modification in pitch or not. **B**. Estimation of the discrimination threshold and accuracy by fitting a sigmoid function. **C,D**. Correlations of the compensation ratios with the discrimination threshold (C) and accuracy (D). *r*_s_ shows Spearman’s signed-rank correlation coefficient.

## Discussion

Recent debates on tight links between motor variability and adaptive regulation have been along with the motor exploration hypothesis, with showing practical evidence in songbirds’ vocalization (Tumer and Brainard, 2007; Andalman and Fee, 2009; Sober and Brainard, 2009, 2012; Charlesworth et al., 2011; Kuebrich and Sober, 2015), and in some other motor actions of humans (Wu et al., 2014) or rodents (Dhawale et al., 2019). Here, we provide further evidence for this debate in human vocalizations by demonstrating that the spontaneous FF variability is positively correlated with the rate of compensatory response against FF shift perturbations induced in the auditory feedback (**Fig. 2A**). This result was consistent with a previous study that used sudden FF shifts in the auditory feedback in the middle of vocalization (Scheerer and Jones, 2012), suggesting robustness of the finding against methodological differences. Further analyses showed that the slowly fluctuating components but not the fast components had the greater impact on the compensatory response (**Fig. 2D,E**). In addition, the compensation ratio for the largest shift conditions (+-100 cent) showed a significant decrease comparing to other shifts (**Fig. 4C**), even exhibiting the correlation with the spontaneous variability of slow component (**Fig. 4D**). These findings provide further support for the idea that spontaneously produced motor noise plays a role in generating motor explorations and results in promoting its adaptive regulation, even in vocal production processes.

Our results further indicated that the slow components of the spontaneous variability more contributed to the compensation than the fast fluctuation (**Fig. 2**), and the main component of the compensation response shared the same frequency range of the slow component (**Fig.3**). The fast fluctuation in vocal FF has been recognized as “microtremor” which is an involuntary fluctuation caused by physical/physiological instability (Schoentgen, 2002), suggesting that this component mainly consists of uncontrollable noise sources generated in the peripheral system. Such peripherally derived variability may not be well suited for adaptation-related motor exploration because of its uncontrollable nature (Dhawale et al., 2017). In contrast, it is indicated that the slow component is controllable in the central nervous system because participants increased the amplitude of FF movement in the range of slow component for compensatory responses. Thus, our result is in concordance with the motor exploration hypothesis, suggesting that the spontaneous variability in slow fluctuation contributes to vocal adaptation by generating the motor exploration.

The present results well fit with the idea that variability in motor production contributes to learning by extending such exploration (Faisal et al., 2008; Renart and Machens, 2014; Wu et al., 2014; Dhawale et al., 2017), and provide further generality of this hypothesis in the vocal control. An alternative explanation for the variability-adaptation relationship could be possible based on a factor of the perceptual ability to detect FF changes. A previous study of vocal FF control reported that children who had less sensitive pitch discrimination abilities showed larger compensations in response to sudden induced FF shifts (Heller Murray and Stepp, 2020), suggesting a possible impact of the auditory ability on the compensation ratio. Although, our result of correlation analysis between perception and compensatory response (**Fig. 5**) did not support this idea since they were not significantly correlated. Thus, we here exclude the possibility of influence from auditory abilities, but employ the spontaneous variability as the main factor explaining the individual difference in the compensation ratio.

More generally, our study suggests a shared strategy in vocal adaptation mechanisms among songbirds and humans. Many studies have shown potential parallels in these two species in vocal learning behaviors and their neural circuitries (Doupe and Kuhl, 1999; Kuhl, 2004; Lipkind et al., 2013; Tchernichovski and Marcus, 2014). Our results add further evidence of such parallels at the level of not only behavioral analogues, but also the computation for vocal adaptation. It should be noted that previous songbird studies have focused on variability and adaptation in a trial-by-trial manner where researchers assessed updating changes in vocal acoustics every song renditions (Kao et al., 2005; Tumer and Brainard, 2007; Sober and Brainard, 2012; Kuebrich and Sober, 2015; Tachibana et al., 2017), although several studies have shown the importance of within-trial variability, i.e., FF fluctuations in one vocal element, on vocal adaptations (Charlesworth et al., 2011; Kojima et al., 2018). Our study here demonstrated the relationship between variability and compensatory responses within each trial in human vocalization, while the relationship between the trial-by-trial variability and updating adaptation over trials will be tested in future studies.

## Methods

### Dataset

The dataset used here was originally obtained in our previous study (Xu et al., 2020). The present study analyzed this in different ways to elucidate the relationship between the variability and compensation behavior in vocal control, although the previous study had focused on the influences of perceptual awareness and vocal responses against manipulating different acoustical features in the auditory feedback. The data were obtained from forty university students (20 females; 18-26 years old) without any experience of formal music training. The experiment was approved by the Human Subjects Ethics Committee of Tokyo Metropolitan University.

The experimental procedure was identical as described in the previous study. In brief, participants were asked to produce isolated vowels /a/ or /u/ according to the letter displayed on a computer screen with hearing auditory feedback via headphones. The auditory feedback was modified by a voice processor (Voice Worksplus, TC Helicon Vocal Technologies, Victoria BC, Canada), and feedbacked to participants with masking pink noise. Participants vocalized twice the same vowel for 2 s with 1 s intermission in each trial, and only the second voice was modified in its feedback (**Fig. 1A**). There was a total of 13 conditions for the second vocalization: 6 for spectral shifts, 6 for spectral-envelope shifts, and 1 for no shift as a control condition. In the spectral shift conditions, the voice spectrum was linearly expanded by ±25, ±50, or ±100 cents (100 cents = 1 semitone), resulting in the shift of the fundamental frequency (FF). The spectral-envelope shift conditions expanded only the envelope by ±3, ±6, or ±12 percent without changing FF. There were 10 trials for each of the 13 conditions for each vowel. The order of 260 trials was pseudo-randomized. Note that we only focused on vocal responses in the spectral shift conditions, but the spectral-envelope shift conditions were excluded from the further analyses in this study. We mainly analyzed the dataset for /a/-vowel trials since the compensatory responses for this vowel was clearer than that for /u/ trials (see **Supporting Information**). After vocalization sessions, participants were also asked to detect whether the modifications had been applied to recorded own voices that were feedbacked to them during the vocalization session. In this listening test, voices in two representative trials were played back to each participant. The participant was asked if they could perceive a change in pitch and/or timbre in the second vocalization comparing with the first one. The present study used these responses to assess the participant’s perceptual ability for detecting the presence of FF shifts in the feedbacked voice.

### Preprocessing

The FF of vocal sound was calculated by Praat 6.0 (Boersma and Weenink, 2017). The FF calculation was performed by an adapted auto-correlation method implemented in the Praat (“To Pitch (ac)”), with 10-ms step, 40-ms window, and frequency boundaries between 75 Hz and 600 Hz. The extracted FF traces were converted into cent values that were in logarithmic scale and obtained as follows: 1200 log_2_(*f / f*_*base*_), where *f* is FF in Hz, and *f*_*base*_ is a base frequency (we used 55 Hz for the base).

We preprocessed the obtained dataset in two steps: alignment and refinement, as described below. We firstly aligned the data by time points of vocal onsets. In this process, the vocal onset and offset were detected from the amplitude envelopes (described below) with a threshold of the background level + 30 dB. The background level was estimated from silent parts of recordings for each participant. Then, we refined the aligned data by detaching or repairing unstable/misdetected data points as follows. Fragmented data points were connected by filling brief temporal gaps (≤40 ms) and removing short fragments (≤50 ms). Unrealistic frequency jumps that were larger than ±100 cents at the beginning part of vocalization were searched backward from 200-ms time point to the onset, and removed. Similarly, unrealistic jumps for the ending parts were also removed by forwardly searching from 300-ms before the offset with the same threshold (±100 cents). After these removals of unstable onset parts, we re-define new onset times as the beginning point of stable vocalization since these unstable data reflected harsh or aperiodic glottal pulsation in which participants could not sense FF shifts in the feedback. Additionally, we also repaired the unrealistic jumps at the middle part of vocalization between 210 to 1500 ms from the vocal onset (filled with the value obtained immediately before the jump).

### Compensation ratio

To quantify compensatory responses against artificial FF shifts in the auditory feedback, we first removed participant-specific frequency changes that were unrelated to the response to FF shifts. For this, a common trend in all trajectories for each participant was removed by subtracting the grand mean of all trials. Moreover, we set the beginning part of each vocalization as zero by subtracting the mean value within a range of 50–150 ms in each trial to measure only the responses to FF shifts. We defined this subtraction baseline period by visual inspection of outcomes of the grand averaging, and excluded the first 50 ms because of its instability. Then, we calculated the mean value of the late part (800–1200 ms) of data, in which the trajectories fluctuated less and were relatively stable (shown as a black bar in **Fig. 1B**). We defined the compensation ratio to quantify the ratio how much the participant compensated own vocal FF against induced FF shifts. This ratio was calculated as a sign-inverted slope of a line (linear regression) fitted to the mean amounts of vocal responses as a function of FF shifts (**Fig. 1C**).

### Variability assessment

To quantify the motor variability in vocalization, we calculated the standard deviation (SD) of the FF within a period between 100 and 1200 ms after the voice onset. For this calculation, we collected FF trajectory data of the first vocalization of each trial, in which no FF shift was presented. We excluded data from trials that followed immediately after the spectral-shifted (and thus FF-shifted) trials to avoid contaminations of possible aftereffects. The computed SDs were averaged for each participant to obtain a variability index from the original (or “whole”) FF trajectories. Then, we computed the mean SD after filtering by a low-pass filter with 5-Hz cutoff, or a band-pass filter with 6–30-Hz bandwidth (second-order Butterworth filter) to obtain the variability index for a slowly fluctuating component (“slow”) or fast fluctuating one (“fast”), respectively. These two frequency bands were defined by visual inspection of the modulation spectrum (**Fig. 2B**). Before filtering, each trajectory was zero-centered by subtracting the mean value to remove the constant component, and filled missing data points with zero. We used the zero-phase digital filtering implemented in MATLAB software (“filtfilt” function).

### Modulation spectrum analysis

For assessing a relative amplitude across different modulation frequencies, we calculated the modulation spectrum by a half-octave-band filter bank. We first upsampled each FF trajectory into a double rate (200 Hz), then, performed centering by subtracting the mean value of it, and filled missing data points with zero. We defined the filter bank as a set of multiple band-pass filters that had 1/2-octave bandwidths with center frequencies equally spaced at 1/4-octave step from 0.4 to 50 Hz (second-order Butterworth filter). The amplitude of each subband was calculated as the root-mean-square value of the filtered trajectory.

### Amplitude calculation

The amplitude envelope of each vocalization was calculated as the root-mean-square (RMS) values of an A-weighted waveform within 40-ms Hanning window for every 10-ms time step by MATLAB software. The obtained amplitude envelope was converted into a logarithmic scale (dB) by a formula: 20 log_10_ (*x*). We calculated the average value of the log-converted amplitude within a period (150-1200ms) that includes the very beginning part of the compensatory response and the plateau part of vocalization. Then, relative values were calculated by subtracting an overall average from all participants’ data.

### Discrimination performance

We quantified the participant’s perceptual ability to detect shifts in own vocal FF using the dataset obtained from the listening test performed after vocalization sessions. We pooled trials irrespective of FF shift directions (minus or plus), and two vowels (/a/ and /u/) to increase the resolution and obtained 8 repetitions (2 directions × 2 vowels × 2 trials) for each of absolute amounts of FF shifts. The detection rate for each absolute FF shift was approximated by fitting a sigmoid function. For this fitting, we used a cumulative probability density function of the normal distribution as the sigmoid. The absolute shift value at 50 % detection rate and the shallowness of fitted sigmoid, which were corresponding to the mean and standard deviation of the cumulative normal distribution, were defined as the discrimination threshold and accuracy, respectively (**Fig. 3B**).

## Acknowledgement

We thank to Drs. Satoshi Kojima, Kouta Kanno, Kentaro Ono for valuable comments on the earlier version of this manuscript. This study was supported by Adolescent Mind & Self-Regulation, Grant-in-Aid for Scientific Research on Innovative Areas, MEXT, Japan (#23118003; Adolescent Mind & Self-Regulation) to R.H. and F.H., Grant-in-Aid for Scientific Research on Innovative Areas, MEXT, Japan (#4903; Evolinguistics) to K.O., MEXT/JSPS KAKENHI Grant No. 16H06525 to F.H., 16H06395 and 16H06396 to R.H., and JSPS Postdoctoral Fellowship, Japan (#269362) to R.O.T.

## Supporting Information

### Vowel difference

**Figure S1.**
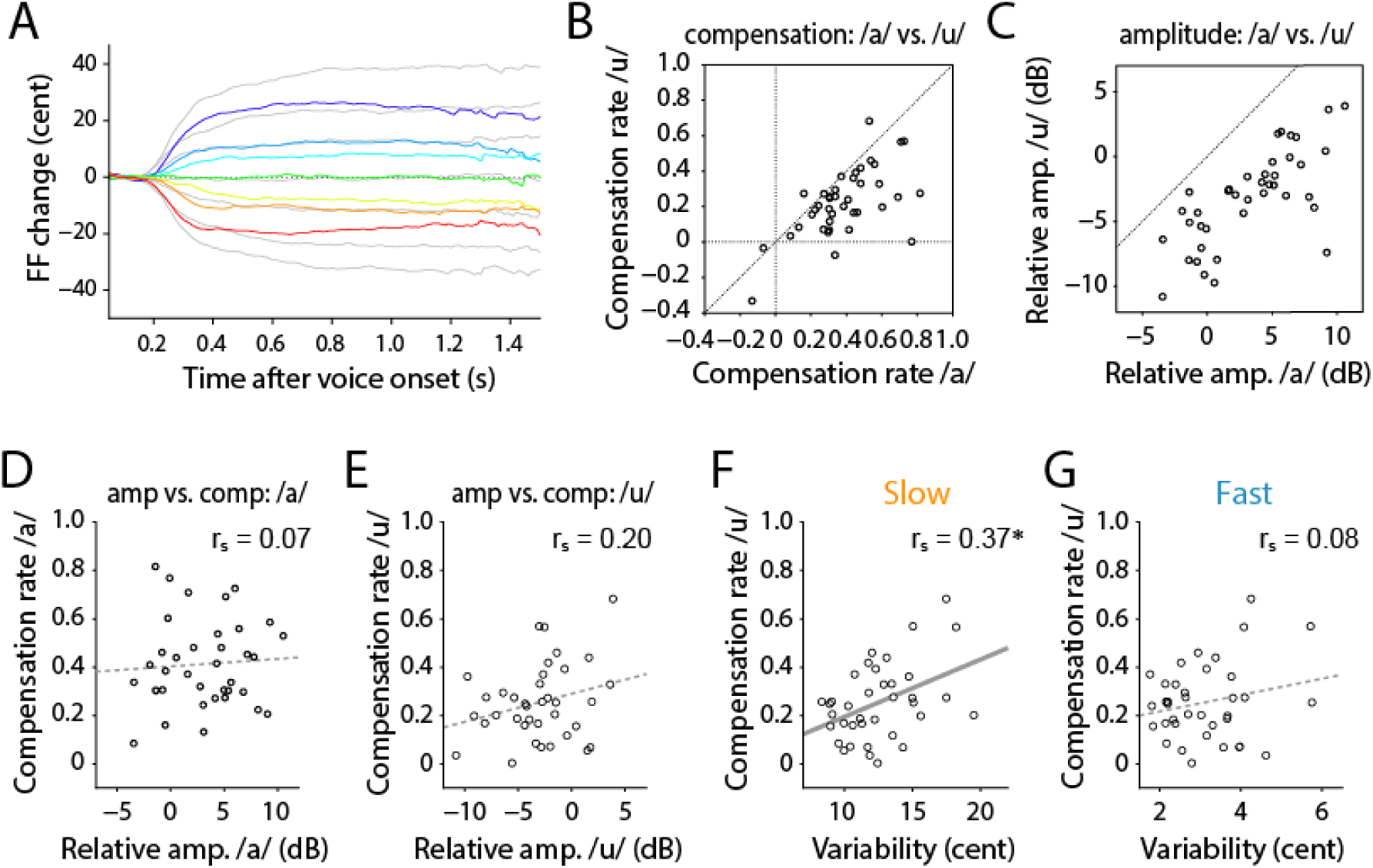
Vowel differences in compensation ratio, amplitude, and variability. **A**. Vocal responses against FF shifts in auditory feedback for /u/ vocalization (colored lines), showing that for /a/ trials as comparisons (gray lines). **B**. The compensation ratio for /u/ vocalizations was generally less than that for /a/. **C**. The voice amplitude of /u/ vowel was generally less than that of /a/. **D,E**. Voice amplitudes of /a/ (D) and /u/ (E) vowels did not show significant correlations with the compensation ratio. **F,G**. Correlation between the compensation ratio and variability of slow (F) or fast (G) components, respectively, for /u/ vocalizations. Asterisk (*) indicates significant correlation (*p* < 0.05).

## Reference

Akagi M, Iwaki M, Minakawa T (1998) Fundamental frequency fluctuation in continuous vowel utterance and its perception. In: 5th International Conference on Spoken Language Processing, pp 1519–1522 Available at: https://www.isca-speech.org/archive/icslp_1998/i98_0027.html.

Akagi M, Kitakaze H (2000) Perception of synthesized singing voices with fine fluctuations in their fundamental frequency contours. In: 6th International Conference on Spoken Language Processing, pp 458–461 Available at: https://www.isca-speech.org/archive/icslp_2000/i00_3458.html.

Andalman AS, Fee MS (2009) A basal ganglia-forebrain circuit in the songbird biases motor output to avoid vocal errors. Proc Natl Acad Sci U S A 106:12518–12523 Available at: http://www.pubmedcentral.nih.gov/articlerender.fcgi?artid=2709669&tool=pmcentrez&rendertype=abstract.

Boersma P, Weenink D (2017) Praat: doing phonetics by computer. Available at: http://www.praat.org/.

Burnett TA, Freedland MB, Larson CR, Hain TC (1998) Voice F0 responses to manipulations in pitch feedback. J Acoust Soc Am 103:3153–3161 Available at: http://www.ncbi.nlm.nih.gov/pubmed/9637026 [Accessed April 20, 2014].

Charlesworth JD, Tumer EC, Warren TL, Brainard MS (2011) Learning the microstructure of successful behavior. Nat Neurosci 14:373–380 Available at: http://dx.doi.org/10.1038/nn.2748 [Accessed March 23, 2014].

Dhawale AK, Miyamoto YR, Smith MA, Ölveczky BP (2019) Adaptive Regulation of Motor Variability. Curr Biol 29:3551-3562.e7 Available at: https://linkinghub.elsevier.com/retrieve/pii/S0960982219311029.

Dhawale AK, Smith MA, Ölveczky BP (2017) The Role of Variability in Motor Learning. Annu Rev Neurosci 40:479–498 Available at: http://www.annualreviews.org/doi/10.1146/annurev-neuro-072116-031548.

Doupe AJ, Kuhl PK (1999) Birdsong and human speech: common themes and mechanisms. Annu Rev Neurosci 22:567–631 Available at: http://www.ncbi.nlm.nih.gov/pubmed/10202549 [Accessed June 11, 2014].

Elman JL (1981) Effects of frequency-shifted feedback on the pitch of vocal productions. J Acoust Soc Am 70:45 Available at: http://www.ncbi.nlm.nih.gov/pubmed/7264071 [Accessed April 26, 2016].

Faisal AA, Selen LPJ, Wolpert DM (2008) Noise in the nervous system. Nat Rev Neurosci 9:292–303 Available at: http://www.nature.com/articles/nrn2258.

Hahnloser RHR, Narula G (2017) A Bayesian Account of Vocal Adaptation to Pitch-Shifted Auditory Feedback. Vasilaki E, ed. PLoS One 12:e0169795.Available at: http://dx.plos.org/10.1371/journal.pone.0169795 [Accessed March 29, 2017].

Hain TC, Burnett T a, Kiran S, Larson CR, Singh S, Kenney MK (2000) Instructing subjects to make a voluntary response reveals the presence of two components to the audio-vocal reflex. Exp brain Res 130:133–141 Available at: http://www.ncbi.nlm.nih.gov/pubmed/10672466.

Hampton CM, Sakata JT, Brainard MS (2009) An avian basal ganglia-forebrain circuit contributes differentially to syllable versus sequence variability of adult Bengalese finch song. J Neurophysiol 101:3235–3245 Available at: http://www.pubmedcentral.nih.gov/articlerender.fcgi?artid=2694116&tool=pmcentrez&rendertype=abstract [Accessed March 23, 2014].

Heller Murray ES, Stepp CE (2020) Relationships between vocal pitch perception and production: a developmental perspective. Sci Rep 10:3912 Available at: http://www.nature.com/articles/s41598-020-60756-2.

Howes P, Callaghan J, Davis P, Kenny D, Thorpe W (2004) The relationship between measured vibrato characteristics and perception in Western operatic singing. J Voice 18:216–230 Available at: https://linkinghub.elsevier.com/retrieve/pii/S0892199703001425.

Kao MH, Brainard MS (2006) Lesions of an avian basal ganglia circuit prevent context-dependent changes to song variability. J Neurophysiol 96:1441–1455 Available at: http://www.ncbi.nlm.nih.gov/pubmed/16723412 [Accessed March 23, 2014].

Kao MH, Doupe AJ, Brainard MS (2005) Contributions of an avian basal ganglia-forebrain circuit to real-time modulation of song. Nature 433:638–643 Available at: http://www.ncbi.nlm.nih.gov/pubmed/15703748.

Kawahara H (1994) Interactions between speech production and perception under auditory feedback perturbations on fundamental frequencies. J Acoust Soc Japan 15:201–202 Available at: https://www.jstage.jst.go.jp/article/ast1980/15/3/15_3_201/_article [Accessed April 26, 2016].

Kojima S, Kao MH, Doupe AJ, Brainard MS (2018) The avian basal ganglia are a source of rapid behavioral variation that enables vocal motor exploration. J Neurosci.

Kuebrich BD, Sober SJ (2015) Variations on a theme: Songbirds, variability, and sensorimotor error correction. Neuroscience 296:48–54 Available at: http://www.ncbi.nlm.nih.gov/pubmed/25305664 [Accessed November 8, 2014].

Kuhl PK (2004) Early language acquisition: cracking the speech code. Nat Rev Neurosci 5:831–843 Available at: http://www.ncbi.nlm.nih.gov/pubmed/15496861 [Accessed June 11, 2011].

Larson CR, Burnett T a, Kiran S, Hain TC (2000) Effects of pitch-shift velocity on voice Fo responses. J Acoust Soc Am 107:559–564 Available at: http://www.ncbi.nlm.nih.gov/pubmed/10641664.

Lipkind D, Marcus GF, Bemis DK, Sasahara K, Jacoby N, Takahasi M, Suzuki K, Feher O, Ravbar P, Okanoya K, Tchernichovski O (2013) Stepwise acquisition of vocal combinatorial capacity in songbirds and human infants. Nature 498:104–108 Available at: http://www.pubmedcentral.nih.gov/articlerender.fcgi?artid=3676428&tool=pmcentrez&rendertype=abstract [Accessed October 8, 2014].

Liu H, Larson CR (2007) Effects of perturbation magnitude and voice F0 level on the pitch-shift reflex. J Acoust Soc Am 122:3671–3677.

Liu P, Chen Z, Larson CR, Huang D, Liu H (2010) Auditory feedback control of voice fundamental frequency in school children. J Acoust Soc Am 128:1306–1312 Available at: http://www.ncbi.nlm.nih.gov/pubmed/20815465 [Accessed November 8, 2014].

Olveczky BP, Gardner TJ (2011) A bird’s eye view of neural circuit formation. Curr Opin Neurobiol 21:124–131 Available at: http://www.pubmedcentral.nih.gov/articlerender.fcgi?artid=3041870&tool=pmcentrez&rendertype=abstract [Accessed June 24, 2011].

Prather J, Okanoya K, Bolhuis JJ (2017) Brains for birds and babies: Neural parallels between birdsong and speech acquisition. Neurosci Biobehav Rev Available at: http://www.ncbi.nlm.nih.gov/pubmed/28087242 [Accessed February 28, 2017].

Renart A, Machens CK (2014) Variability in neural activity and behavior. Curr Opin Neurobiol 25C:211–220 Available at: http://www.sciencedirect.com/science/article/pii/S0959438814000488 [Accessed April 29, 2014].

Saitou T, Unoki M, Akagi M (2005) Development of an F0 control model based on F0 dynamic characteristics for singing-voice synthesis. Speech Commun 46:405–417 Available at: http://linkinghub.elsevier.com/retrieve/pii/S0167639305000993.

Scheerer NE, Jones JA (2012) The relationship between vocal accuracy and variability to the level of compensation to altered auditory feedback. Neurosci Lett 529:128–132 Available at: http://www.ncbi.nlm.nih.gov/pubmed/22995182 [Accessed November 8, 2014].

Schoentgen J (2002) Modulation frequency and modulation level owing to vocal microtremor. J Acoust Soc Am 112:690–700 Available at: http://asa.scitation.org/doi/10.1121/1.1492820.

Shipp T, Sundberg J, Doherty ET (1988) The effect of delayed auditory feedback on vocal vibrato. J Voice 2:195–199 Available at: https://linkinghub.elsevier.com/retrieve/pii/S0892199788800766.

Sober SJ, Brainard MS (2009) Adult birdsong is actively maintained by error correction. Nat Neurosci 12:927–931 Available at: http://dx.doi.org/10.1038/nn.2336 [Accessed March 26, 2014].

Sober SJ, Brainard MS (2012) Vocal learning is constrained by the statistics of sensorimotor experience. Proc Natl Acad Sci U S A 109:21099–21103 Available at: http://www.pubmedcentral.nih.gov/articlerender.fcgi?artid=3529072&tool=pmcentrez&rendertype=abstract [Accessed March 23, 2014].

Sundberg J (1987) The Science of the Singing Voice. Northern Illinois University Press. Available at: https://books.google.co.jp/books?id=iYGNQgAACAAJ.

Tachibana RO, Takahasi M, Hessler NA, Okanoya K (2017) Maturation-dependent control of vocal temporal plasticity in a songbird. Dev Neurobiol 77:995–1006 Available at: http://doi.wiley.com/10.1002/dneu.22487 [Accessed February 13, 2017].

Tchernichovski O, Marcus G (2014) Vocal learning beyond imitation: Mechanisms of adaptive vocal development in songbirds and human infants. Curr Opin Neurobiol 28:42–47 Available at: http://dx.doi.org/10.1016/j.conb.2014.06.002.

Tumer EC, Brainard MS (2007) Performance variability enables adaptive plasticity of “crystallized” adult birdsong. Nature 450:1240–1244 Available at: http://www.ncbi.nlm.nih.gov/pubmed/18097411 [Accessed March 23, 2014].

Woolley SC, Kao MH (2015) Variability in action: Contributions of a songbird cortical-basal ganglia circuit to vocal motor learning and control. Neuroscience 296:39–47 Available at: http://www.ncbi.nlm.nih.gov/pubmed/25445191 [Accessed December 8, 2014].

Wu HG, Miyamoto YR, Gonzalez Castro LN, Ölveczky BP, Smith MA (2014) Temporal structure of motor variability is dynamically regulated and predicts motor learning ability. Nat Neurosci 17:312–321 Available at: http://www.pubmedcentral.nih.gov/articlerender.fcgi?artid=PMC4442489 [Accessed July 22, 2014].

Xu M, Tachibana RO, Okanoy K, Hagiwara H, Hashimoto R, Homae F (2020) Unconscious and distinctive control of vocal pitch and timbre during altered auditory feedback. Front Psychol.

